# Cytotoxic efficiency of human CD8^+^ T cell memory subtypes

**DOI:** 10.1101/2021.03.15.435339

**Authors:** Arne Knörck, Gertrud Schwär, Dalia Alansary, Lorenz Thurner, Markus Hoth, Eva C. Schwarz

## Abstract

Immunological memory is an important concept to protect humans against recurring diseases. Memory CD8^+^ T cells are required for quick expansion into effector cells but also provide immediate cytotoxicity against their targets. Whereas many functions of the two main cytotoxic subtypes, effector memory CD8^+^ T cells (T_EM_) and central memory CD8^+^ T cells (T_CM_), are relatively well defined, single T_EM_ and T_CM_ cell cytotoxicity has not been quantified. Here, we analyze human CD8^+^ subtype distribution following SEA stimulation and quantify the expression of death mediators, including perforin, granzyme B, FasL and TRAIL. We find higher perforin, granzyme B and FasL expression in T_EM_ and compared to T_CM_. To quantify single T_EM_ and T_CM_ cytotoxicity, we develop a FRET-based fluorescent assay with NALM6 target cells stably transfected with a GFP-RFP FRET construct based on a caspase-cleavage sequence (apoptosis sensor Casper-GR). Applying this assay, T_EM_ or T_CM_ induced target cell apoptosis or necrosis can be quantified. We find that single T_EM_ are much more effective than T_CM_ in killing their targets mainly by apoptosis and secondary necrosis. The reason for this is the higher perforin expression and on a more efficient lytic immunological synapse during T_EM_-target contact compared to T_CM_-target contact. Defining and quantifying single T_EM_ and T_CM_ cytotoxicity and the respective mechanism should be helpful to optimize future subset-based immune therapies.

## Introduction

Cellular cytotoxicity mediated by CD8^+^ activated cytotoxic T cells (CTL) is a crucial function of the adaptive immune system to efficiently fight cancer or viral infections. CTL kill virus-infected cells or tumor cells by various effector mechanisms. Directed release of lytic granules containing perforin and granzymes or death receptor engagement mediated by FasL or TRAIL are the most prominent mechanisms (1–5).

CTL are central of modern cancer immunotherapy. In the ideal scenario, CTL specifically detect MHCI-presented antigens on the surface of tumor cells and finally kill these targets (6, 7). The outcome of the therapy using genetically modified T cell populations is influenced by central or peripheral tolerance, tumor- or virus-associated immune suppression, functional defects of the MHCI-machinery mitigating presentation of tumor neo- or viral antigens or the occurrence of severe side-effects like the cytokine release syndrome (CRS) or neurotoxicity (NT) (8, 9). To date, the majority of current T cell therapies rely not on defined T cell subsets but the entire T cell population (10). However, recent data support the idea that pre-selection of T cell subsets might help to optimize further treatments (11–13). Unfortunately, it is problematic to achieve this because patients are often already very sick with severe lymphopenia. Nevertheless, a detailed understanding CTL subset cytotoxicity is needed to optimize future cellular T cell therapies.

T cell memory subsets, T_EM_ or T_CM_, differentiate during an immune response and provide long term immunity against recurring viral or tumour antigens. The different subpopulations are widely defined by the surface expression of CCR7/CD62L and CD45RO/RA (14, 15). Whereas T_EM_ likely respond immediately to a recurring antigen, T_CM_ reside in the secondary lymphatic tissues keeping a comparably high proliferative capacity to develop a new population of effector cells after a second conjugation to the same antigen (16, 17). Although T_EM_ are expected to have a higher cytotoxic potential compared to T_CM_, detailed functional information about the orchestration of different killing mechanisms and the killing machinery which is used by distinct subtypes, is still missing.

In this study, we developed a single cell kinetic assay to quantify cancer cell apoptosis and necrosis following T_EM_ or T_CM_ contact. We show that T_EM_ and T_CM_ utilized both, perforin and death receptor mediated mechanisms to kill target cells. While perforin-mediated induction of apoptosis is the most prevalent mechanism for both subsets, T_EM_ displayed an increased cytotoxic potential that is not only caused by higher perforin expression but we also find a shortened time to establish the first killer-target cell contact with a subsequent successful killing.

## Results

### Subtype distribution of SEA-stimulated CD8^+^ T cells (SEA-CTL)

Upon activation, cytotoxic T-lymphocytes (CTL) differentiate into various subsets to execute their cytotoxic function. In this study, we used a widely accepted protocol for activation of T cells by staphylococcal enterotoxin A (SEA). PMBC were isolated from leukoreduction system (LRS) chambers and pulsed with SEA. CTL were positively isolated after the expansion of the mixed PBMC population for 5 days (Fig. 1A).

**Figure 1:**
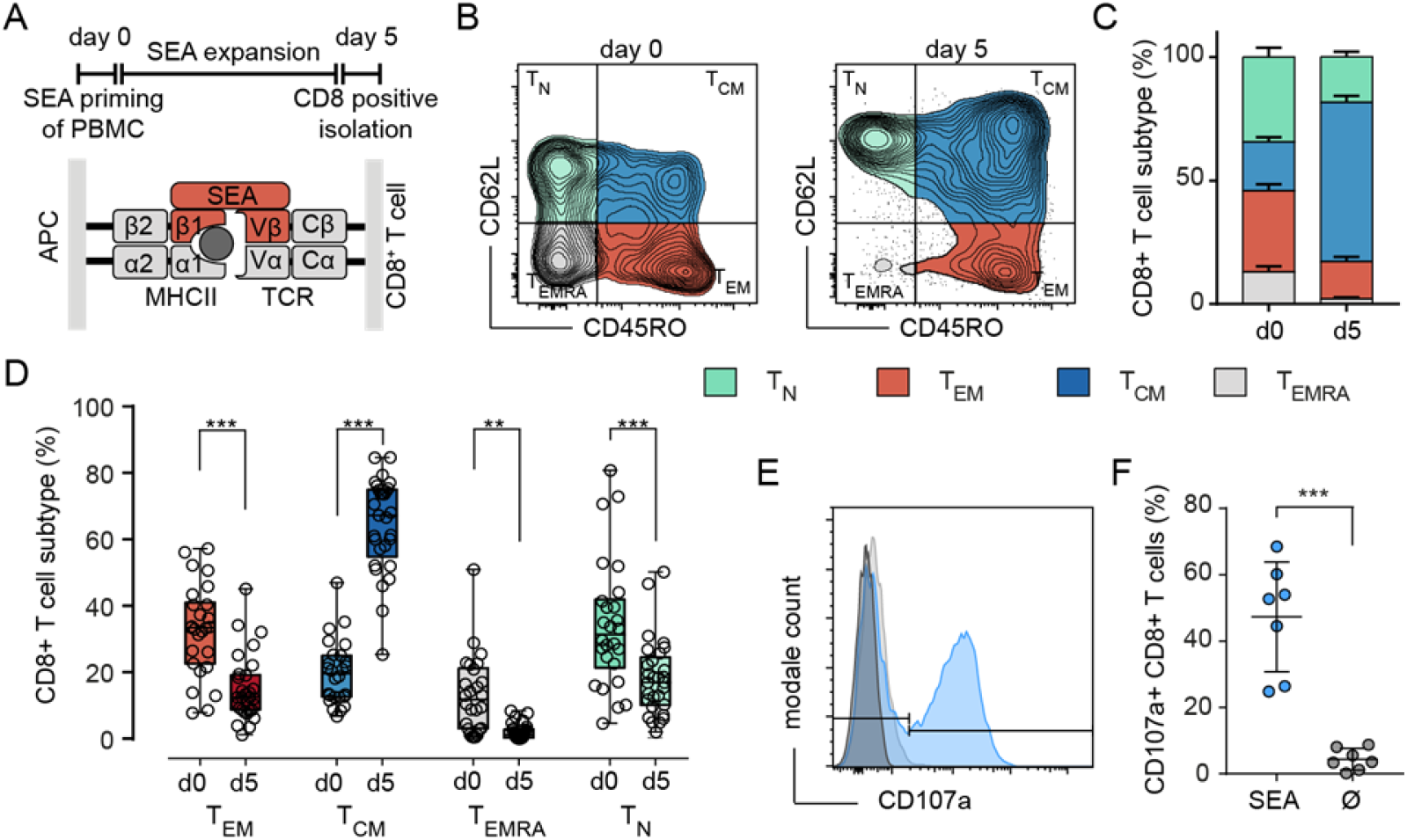
SEA stimulation induces differentiation mainly into a T_CM_ subtype. A) Scheme of the 5-day stimulation with SEA (Staphylococcal enterotoxin A) and the mechanism of CTL-target cell crosslinking by the SEA molecule. B) Representative example for gating of (CD3^+^/CD8^+^) CTL into T_N_ (CD62L^+^/CD45RO^-^), T_CM_ (CD62L^+^/CD45RO^+^), T_EM_ (CD62L^-^/CD45RO^+^) and T_EMRA_ (CD62L^-^/CD45RO^-^) on day 0 and day 5 of SEA-stimulation. C), D) n= 26-29 donors. C) Quantification of subtype distribution at day 0 and day 5. Data are presented as mean+SEM. D) Compared frequencies of T_N_, T_CM_, T_EM_ and T_EMRA_ on day 0 and day 5. Box and Whisker plots show min, max, median and 25-75% interquartile range. Statistical analysis was done by a two-way ANOVA. E, F) Detection of SEA-dependent degranulation by CD107a surface expression of SEA-CTL after co-incubation with SEA-pulsed Raji cells. E) Degranulation of cells from a representative donor (−SEA: grey, +SEA: blue. F) Quantification of (E) with SEA-loaded (SEA) and control (no SEA, Ø) target cells. Data are shown as mean±SD of 7 donors.

CD8^+^ naïve and memory T cell subsets were identified by the surface molecules CD62L or CCR7 and CD45 isoforms RA and RO (15). Fig. 1B illustrates the differentiation of the CD8^+^ population towards a T_CM_ phenotype following SEA stimulation. The frequency of the T_CM_ subset increased from 19.8 ±9.5% on day 0 to be the predominant subset (64±5%) on day 5 of SEA-stimulation. Frequencies of T_EM_ and T_N_ were decreased by roughly 50% (T_EM_ 53.9%, T_N_ 46.7%) to 15.1% and 18.3% of the CD8^+^ T cell population, while most of the T_EMRA_ cells disappeared during the expansion period (Fig. 1C). Healthy human donors show a huge variation in subtype distribution but T_CM_ was always the prominent subtype after SEA stimulation whereas T_EM_ and T_N_ decreased, and T_EMRA_ was almost absent (Fig. 1D). To test the SEA reactivity of stimulated CTL (SEA-CTL), we analyzed the degranulation after co-incubation with SEA-pulsed target cells by detection of surface-located CD107a (Fig. 1E, F). SEA-dependent degranulation shows a wide spread of donor-dependent variation. On average, 47.3±16.5% of SEA-CTL degranulated in case target cells were loaded with SEA compared to 4.4±3.3% in case target cells were not loaded with SEA (Fig. 1F).

To assess the cytotoxic potential of each individual subset, we determined the perforin expression in T_N_, T_CM_, T_EM_ and T_EMRA_ using two different anti-perforin antibody clones (Fig. 2A). The antibody clone δG9v identifies granule-associated perforin ready to lyse the target cell whereas clone B-D48 detects also newly synthesized perforin thus reflecting the total perforin content of the cell {Hersperger, 2008 #12). The overall and the lytic granule-associated content of perforin were higher in T_EM_ compared to T_CM_. Although the frequencies of perforin expressing cells are comparable among T_EM_ (70.17±12%) and T_CM_ (65.06±8.9%) (Fig. 2A), the analysis of the perforin MFI-ratios reveals that T_EM_ express 2.8 times more granular and 2.0 times more total perforin than T_CM_ (Fig. 2B). In addition, the death receptors FasL and TRAIL could be detected on the surface of T_CM_ and T_EM_. However, only FasL (Fig. 2C, D) but not TRAIL (Fig. 2E) were upregulated after SEA-loaded target cell contact. T_EMRA_ express a significantly increased amount of granular and overall perforin (Fig. 2B) but the subpopulation is almost absent after SEA-stimulation (2.14±2.4% of all cells). In conclusion, after expansion by SEA, the SEA-CTL population consists of mainly cytotoxic-competent T_EM_ and T_CM_ against SEA-loaded target cells.

**Figure 2:**
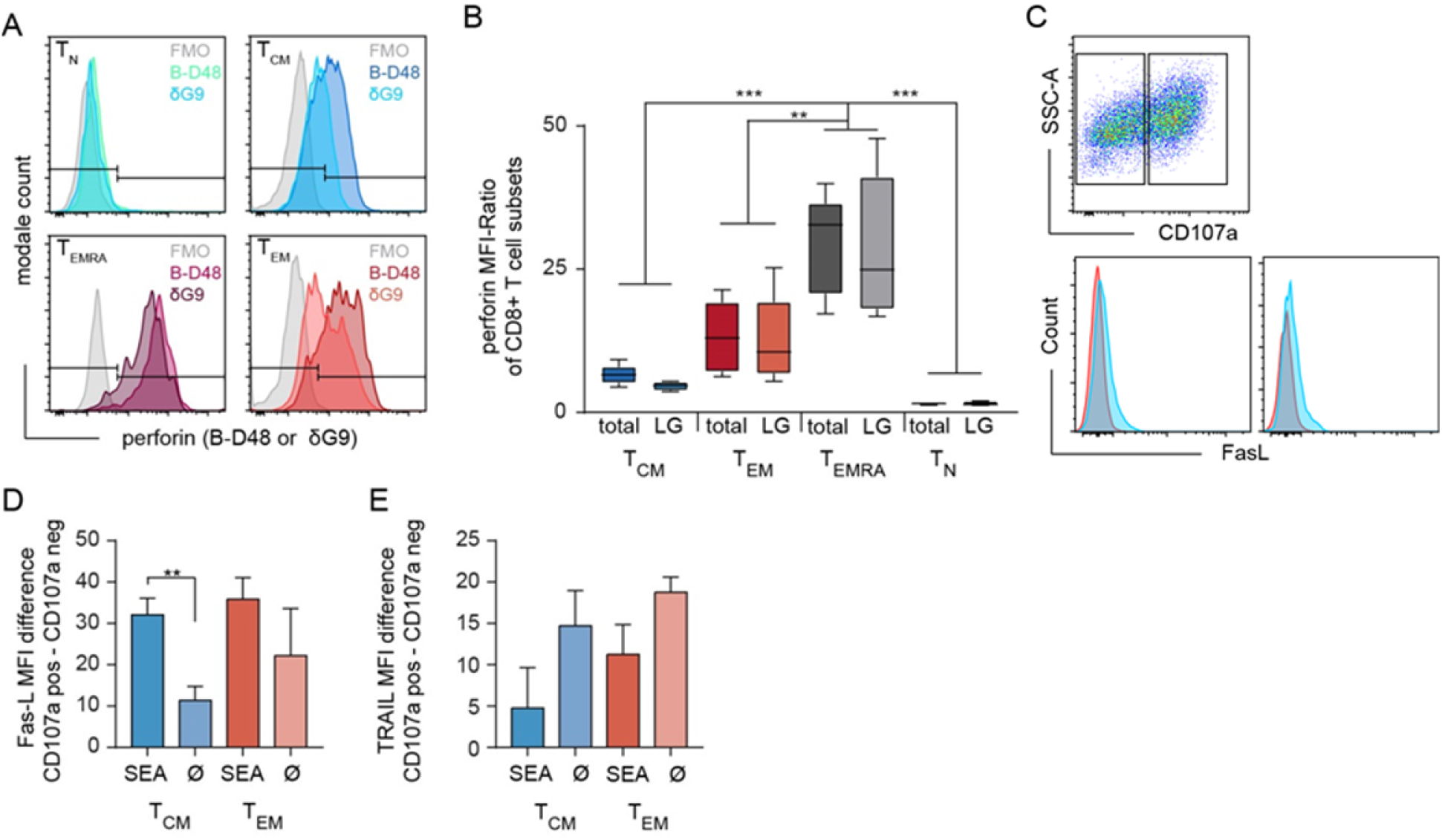
Expression of death mediators in SEA-CTL. A) Content of intracellular perforin stores analyzed in SEA-CTL (bead-isolated SEA-stimulated CD8^+^ population) subsets defined by CCR7 and CD45RO (clone B-D48 for total perforin, clone δG9 for perforin in lytic granules) shown for a representative donor. B) Quantification of subset specific perforin in lytic granules (LG, δG9) or overall (total, B-D48) by MFI-Ratio (MFI Perforin^+^ Population/MFI FMO) shown for 5 donors. Box and Whisker plots show min, max, median and 25-75% interquartile range. Statistical analysis was done by a two-way ANOVA. C) CTL-SEA were co-incubated with 3}10^5^ SEA-pulsed NALM6 at an E:T ratio of 2:1 for 4h in presence of CD107a antibody. Cells were collected and stained with CD45RA, CCR7, CD95L and TRAIL (CD253) antibodies. CD107a^+^ and CD107a^-^ cells were gated for each subset and FasL expression was analyzed. Quantification of FasL (D) or TRAIL expression on degranulating cells (E) on SEA-CTL (SEA) or control cells (Ø) in T_CM_ and T_EM_ subpopulations. Data are shown as mean+SD, n=3.

### Subset specific cytotoxic potential of T_EM_ and T_CM_ subpopulations

To determine the cytotoxic potential of individual memory subsets we sorted bead-isolated SEA-stimulated CD8^+^ population (SEA-CTL) into CD62L^+^/CD45RO^+^ T_CM_ and CD62L^-^/CD45RO^+^ T_EM_ subpopulations (Fig. 3A, pre- and post-sort). We also sorted CD62L^+^/CD45RO^-^ T_N_ from SEA-CTL of 4 different donors. We used these T_N_ as an internal negative control, although the cells cannot be considered as truly naïve since they were in the environment of SEA stimulation for 4 days. However, in terms of their surface markers, they are considered T_N_. 24h after subpopulation sorting we re-analyzed the sorted T_EM_ or T_CM_ cells and found a purity of >80% (Fig. 3B). At the same time, we analyzed the killing capacity of the different subpopulations in a time-resolved population killing assay (Fig. 3C). This assay allows quantification of the cytotoxic potential at different time points and quantification of the maximum killing rate which describes the maximal difference in target cell lysis between two measurement points (10 min). As expected, T_N_ basically did not kill target cells. The T_EM_ subpopulation displayed the highest killing efficiency as quantified by the maximal killing rate (Fig. 3D) and the target cell lysis at 120 min (Fig. 3E) compared to T_CM_, T_N_ and the entire SEA-CTL population as control. Interestingly, the change of target lysis was not different during the last 2 hours (between 120 min and 240 min) among T_EM_, T_CM_ and SEA-CTL (Fig. 3F, compare also Fig. 3C).

**Figure 3:**
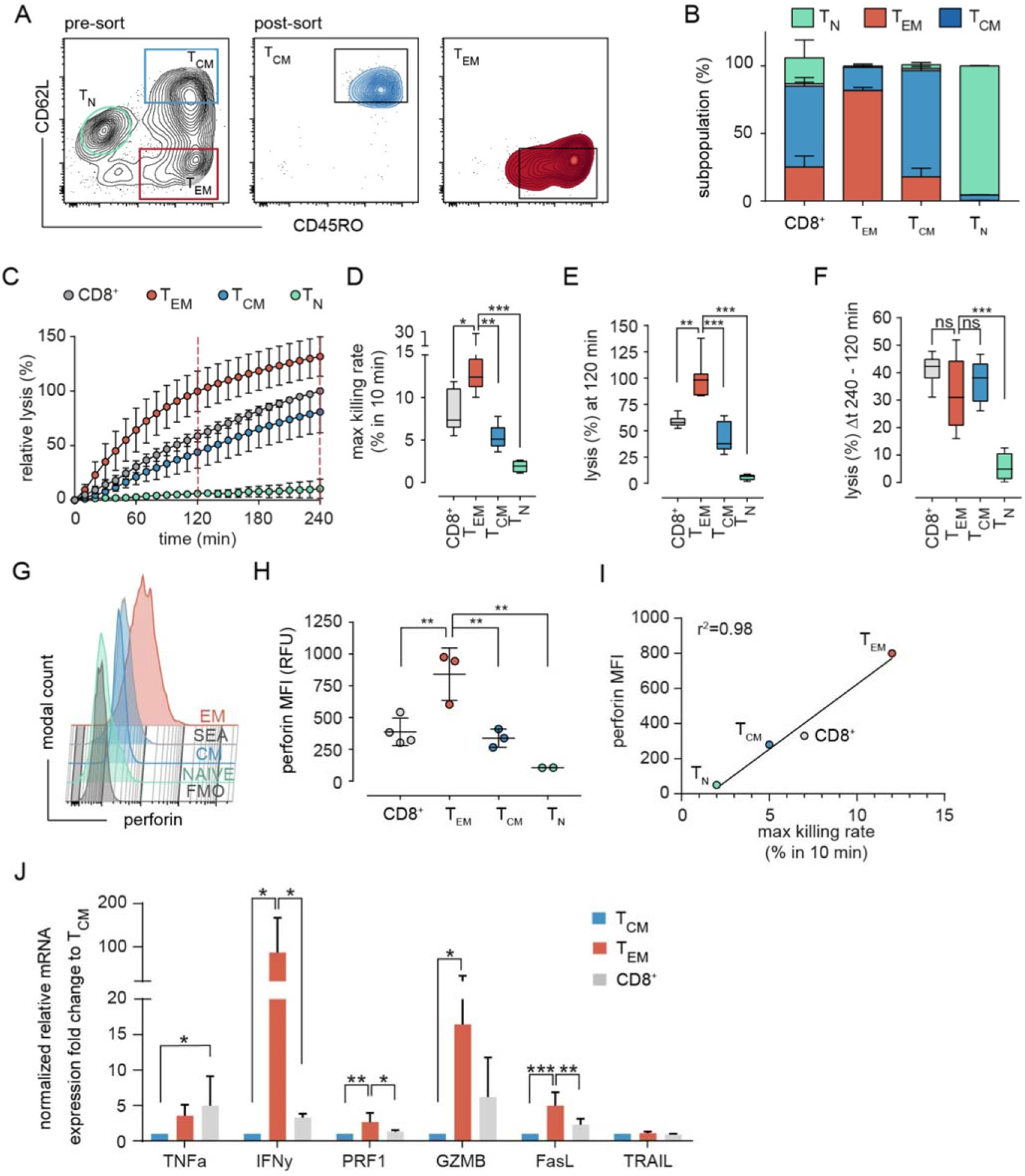
Subset-specific cytotoxicity of sorted T_EM_ and T_CM_. A) Representative example of the sorting strategy. After exclusion of doublets and dead cells by light scatter parameters, the SEA-CTL (CD8^+^) population was separated and sorted into T_EM_ and T_CM_ cells by the surface marker CD62L and CD45RO. 24h after sorting cells were re-analyzed by a staining of CD45RA and CCR7. The subset composition of sorted samples is shown as stacked bar graphs B), (T_EM_, red; T_CM_, Blue; T_N_ (green). C) Cytotoxicity of sorted T_EM_, T_CM_, T_N_ and of the unsorted SEA-CTL population (CD8^+^) was analyzed by a calcein-based real-time killing assay. D-F) To quantify the killing efficiency, the maximal killing rate (maximal target cell lysis within a 10 min interval), the target lysis between 0 - 120 min (E) and 120 min - 240 min (F) were analyzed (number of donors, SEA-CTL (CD8^+^) n=9, T_EM_ n=8, T_CM_ n=7, T_N_ n=4 donors. Box and Whisker plots in D -F show min, max, median and 25-75 % interquartile range. Statistical analysis was done by a one-way ANOVA. G) Perforin expression in sorted T_EM_, T_CM_, T_N_ and in the unsorted SEA-CTL (CD8^+^) population of a representative donor analyzed by flow cytometry. H) Quantification of intracellular perforin by MFI. SEA-CTL (CD8^+^) n=4, T_EM_ n=3, T_CM_ n=3, T_N_ n=2 donors. Scatter plots show mean±SD, statistical analysis was done by a one-way ANOVA. I) MFI of intracellular perforin (H) plotted against maximal killing rates (D). J) Expression of various effector molecules, TNFa, IFNy, PRF1, GZMB, FASL and TRAIL genes was analyzed on mRNA level by RT-qPCR, 48h after sorting. Relative mRNA expression is shown as fold change normalized to the T_CM_ population. T_EM_ and T_CM_ n=4, SEA-CTL (CD8^+^) n=2 donors. Bar graphs show means ± SD. Statistical analysis was done by a Kruskal-Wallis-Test.

Since perforin-mediated exocytosis is known to be rapidly executed {Hassin, 2011 #18;Li, 2014 #19;Meiraz, 2009 #21;Prager, 2019 #22}, we explored the perforin content in sorted subsets in parallel to the population killing assay to confirm the increased expression of granule-associated perforin (using antibody clone ΔG9), which we observed for T_EM_ before in in non-sorted subsets (Fig. 2A, B). Again, we found a significant difference in perforin content of T_EM_ compared to T_CM_ cells (Fig. 3G, H). The SEA-CTL (CD8^+^) control-population expressed slightly more perforin than T_CM_ which is in good agreement with the subtype composition (T_CM_>T_EM_>T_N_). T_N_ sorted cells have almost no perforin (Fig. 3H) in good accordance with the negligible killing efficiency shown before (Fig. 3C). The mean fluorescence intensity of perforin correlated almost perfectly with the maximal killing rate of the subtypes and the entire SEA-CTL (CD8^+^) population (Fig. 3I, r^2^=0.98). Therefore, it appears very likely that the differences of initial lysis of target cells during the first 120 min are caused by differences in perforin content.

When analyzing various effector molecules important for killing mechanisms on mRNA level, we also found interferon-γ, granzyme B and FasL but not TNFα or TRAIL significantly enhanced in T_EM_ (Fig. 3J), suggesting that additional, possibly slower killing mechanism than rapid lysis by perforin might contribute to the enhanced T_EM_ mediated target cell lysis.

### Analysis of T_EM_ and T_CM_ induced target cell lysis at single cell level using the apoptosis sensor Casper-GR

We assumed that the differences in subset-specific cytotoxicity measured with the population killing assay were most likely caused by perforin-mediated target cell killing (Fig. 3C). However, the population-mediated assays do not allow discriminating apoptotic or necrotic target cell death. Numerous tools to dissect the different mechanisms during target cell lysis have been developed in the past years (18–21) but the diversity between different subtypes was not in the focus of this research. The microscope-based assay recently introduced (18) allows the detailed analysis and identification of apoptotic and necrotic lysis events on a single cell level by using the FRET-based apoptosis sensor Casper-GR (22). Up to now, this assay is solely published for natural killer (NK) cells (18). To establish and validate this assay for human CTL, we first tried to stably transfect the clonal B cell lymphoma target cell line Raji (used for the population killing assay, Fig. 3C) with pCasper-GR. However, expression levels were highly variable compromising a detailed quantification. Therefore, we turned to the target cell line NALM6, a B cell line established from a patient with acute lymphoblastic leukemia (ALL) which can also be loaded with SEA.

To validate NALM6 and Raji cells against each other, we compared CTL cytotoxicity in the population killing assay. Neither the kinetics of killing nor the maximal killing rate were different (Supplementary Fig. 1) indicating that human CTL eliminate both targets equally efficient. Therefore, we used stably-transfected Casper-GR NALM6 cells (from now on named NALM6 pCasper) in the following experiments as target cells. 98.7% of the NALM6 pCasper population was homogenously and strongly positive for GFP and RFP signals (Fig. 4A, B). To monitor the viability of NALM6 pCasper under experimental conditions (37°C, 5% CO_2_) we recorded the GFP and FRET signals at 5 min intervals which remained stable over the complete 8 hours measurement period (Fig. 4C). First, we tested the functionality of the apoptosis sensor without using SEA-CTL effector cells by staurosporine, which induces caspase-3 mediated apoptosis (23). We also tested the induction of death receptor-mediated apoptosis by the anti-Fas antibody Apo1-1 mimicking FasL and by recombinant TRAIL to induce TRAIL-receptor mediated cell death over 10 hours (Fig. 4D). The quantitative analysis of cell lysis is illustrated in a stacked diagram over time called death plots. These illustrate the kinetic distribution of lysed cells (dark grey), apoptotic (green) and living cells (light grey) (Fig. 4D, right) After 10 hours, in comparison to the untreated living control, staurosporine and recombinant TRAIL induced significantly enhanced apoptotic cell deaths (95.7% ± 1.5% and 71.8% ± 12.3%, respectively) whereas Apo1-1 induced only very little apoptotic cell death. Interestingly, by simple counting the number of cells, the calculated growth factor was significantly decreased in Apo1-1 treated cells compared to untreated control cells meaning the anti-Fas antibody reduced the proliferation of NALM6 pCasper (Fig. 4E, right panel).

**Figure 4:**
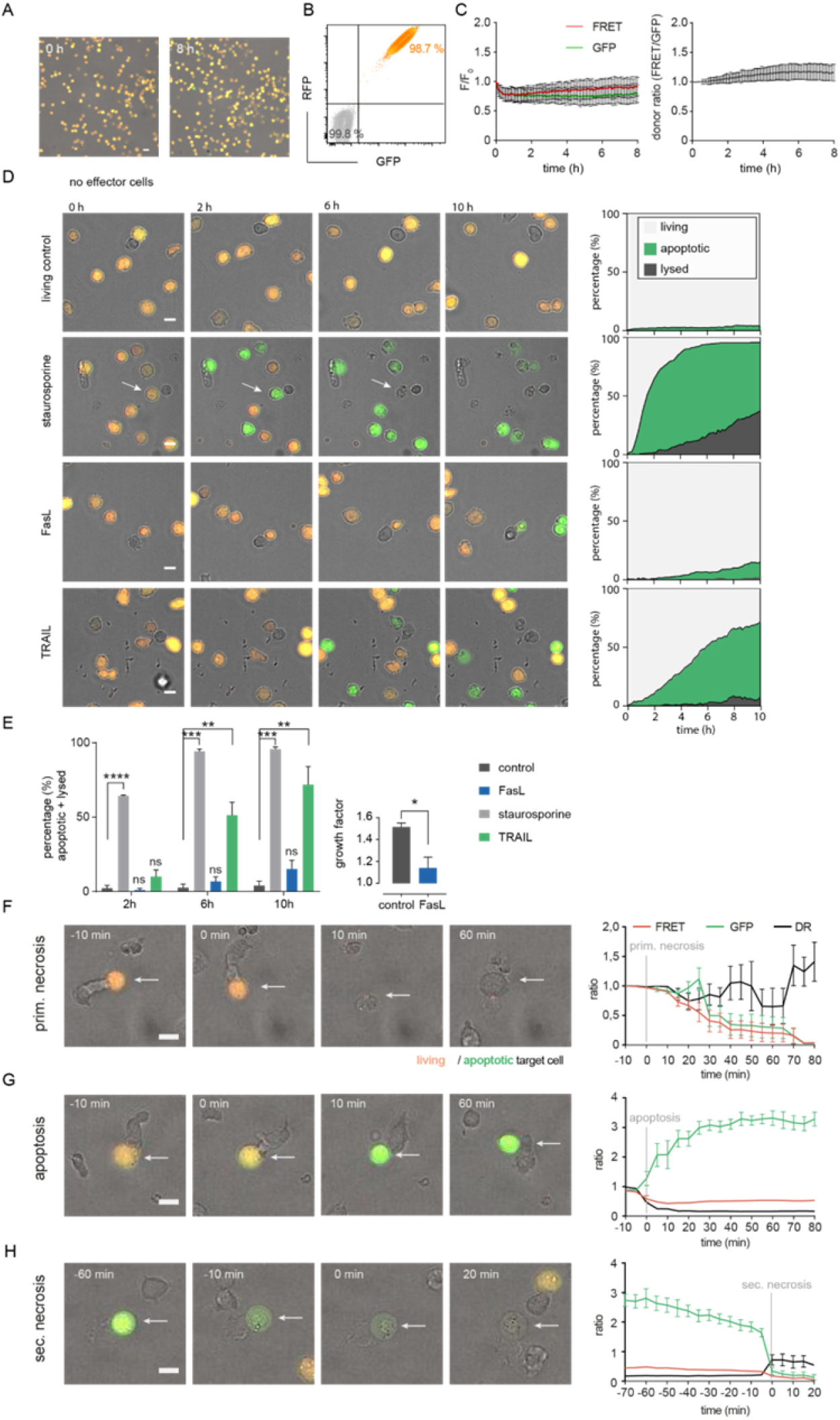
Establishing NALM6 pCasper at single cell level as target cells for SEA-CTL to distinguish between apoptotic and necrotic cell death. A) Viability of NALM6 pCasper cells at 37° C, 5% CO2 for 8h. Viable cells are orange, apoptotic cells are green. B) NALM6 cells were stably transfected with pCasper-GR. GFP and RFP fluorescence of NALM6 pCasper in comparison to non-transfected NALM6 cells (grey) detected by flow cytometry as dot blot. C) Time-resolved averaged normalized GFP and FRET fluorescence (left) and FRET donor ratio (right) over 8h. D,E) Detection of apoptosis induction by staurosporine, anti FasL or recombinant TRAIL. D) Representative images and automated death plot analysis of NALM6 pCasper cells (control, n=426 cells) over 10h treated with 20 μM staurosporine (n=283 cells), 5 μg/ml APO1-1 (n=254 cells) or 5 μg/ml recombinant TRAIL (n=405 cells) are shown. E) Quantification of the fraction of apoptotic and lysed cells and the effect of Apo1-1 treatment (p=0.037) on the growth factor (mean #cells (0-60 min) / #cells (600 min)), single donor, duplicates. Statistical analysis was done using a Friedman test. Specific combinations of substances were used to inhibit perforin and death receptor mediated cytotoxic mechanisms during the assay and by 2h of preincubation. F) Representative lysis event showing a primary necrosis and time-resolved averaged normalized GFP and FRET fluorescence of five cells. G) Representative lysis event showing an apoptosis and time-resolved averaged normalized GFP and FRET fluorescence of seven cells. H) Representative lysis event showing a secondary necrosis following an apoptosis and time-resolved averaged normalized GFP and FRET fluorescence of seven cells. F-H) SEA-CTL were used as effector cells.

Next, we analyzed the modes of cell death occurring in NALM6 pCasper after contact with SEA-CTL. Target cell apoptosis and necrosis following CTL contact could be distinguished as previously established for NK cells (18). We found examples for primary necrosis (Fig. 4F), apoptosis (Fig. 4G) and secondary necrosis (Fig. 4H). Primary necrosis of NALM6 pCasper target cells induced by SEA-CTL is characterized by the parallel loss of the GFP- and FRET fluorescence signals accompanied by morphological changes (Fig. 4F). The induction of caspase-mediated apoptosis following SEA-CTL contact was identified by an increase of the GFP-fluorescence signal and a decrease of the FRET-donor ratio (Fig. 4G). The GFP- and RFP-molecules of the FRET-construct are linked by the caspase recognition sequence DEVD which is cleaved by different caspases after induction of apoptosis (caspase-3, -7 and others). Secondary necrosis following initial apoptosis is characterized by a sudden drop of GFP- and FRET-fluorescence signals accompanied by an increase of the cell volume typically for necrotic cell death (Fig. 4H). Thus, NALM6 pCasper are suitable targets to study different mechanisms of CTL-mediated target cell death.

### Single cell level T_EM_ and T_CM_ analyses reveal distinct cytotoxic efficiencies

The population real-time killing assay revealed clear differences between T_EM_ and T_CM_ cytotoxic efficiencies (Fig. 3C). However, it does not allow any conclusions on mechanisms. Therefore, we used the single cell killing assay to compare T_EM_ and T_CM_ to obtain insights into potential mechanisms. Fig. 5A shows representative examples of individual NALM6 pCasper target cells which are killed after contact with sorted T_EM_ or T_CM_. The latter ones are difficult to see because they are not stained. We therefore depicted an example in the insets to illustrate the contact between T_EM_ or T_CM_ with the targets. Initially all target cells are vital (orange color). The kinetic analysis revealed that T_EM_ and T_CM_ initially mainly induce apoptosis in NALM6 targets as indicated by the green color. We quantified the induction of target cell apoptosis in death plots indicated by the green areas in Fig. 5B. T_EM_ were more efficient to induce apoptosis than T_CM_, which evident from the initial part of the curves and also at the time point of 2 hours (Fig. 5C). Over longer times, both T_EM_ and T_CM_ eliminated almost all targets by apoptosis (Fig. 5B) as quantified at 4 hours (Fig. 5D) and 8 hours (Fig. 5E). The faster apoptosis induction of T_EM_ compared to T_CM_ is also evident from the median time duration of 70 min until a target cell in the T_EM_ data set was apoptotic compared to 180 min in the T_CM_ data set (Fig. 5F).

The death plots (Fig. 5B) also indicate that after initial apoptosis, secondary necrosis (light grey) was induced in many of the targets. Again, T_EM_ were faster to induce secondary necrosis than T_CM_ (Fig. 5B) which is quantified for the different time points of 2, 4, and 8 hours (Fig. 5C-E). Finally, only T_EM_ but not T_CM_, could induce primary necrosis (without preceding apoptosis) as indicated by the dark grey color (Fig. 5B-E).

**Figure 5:**
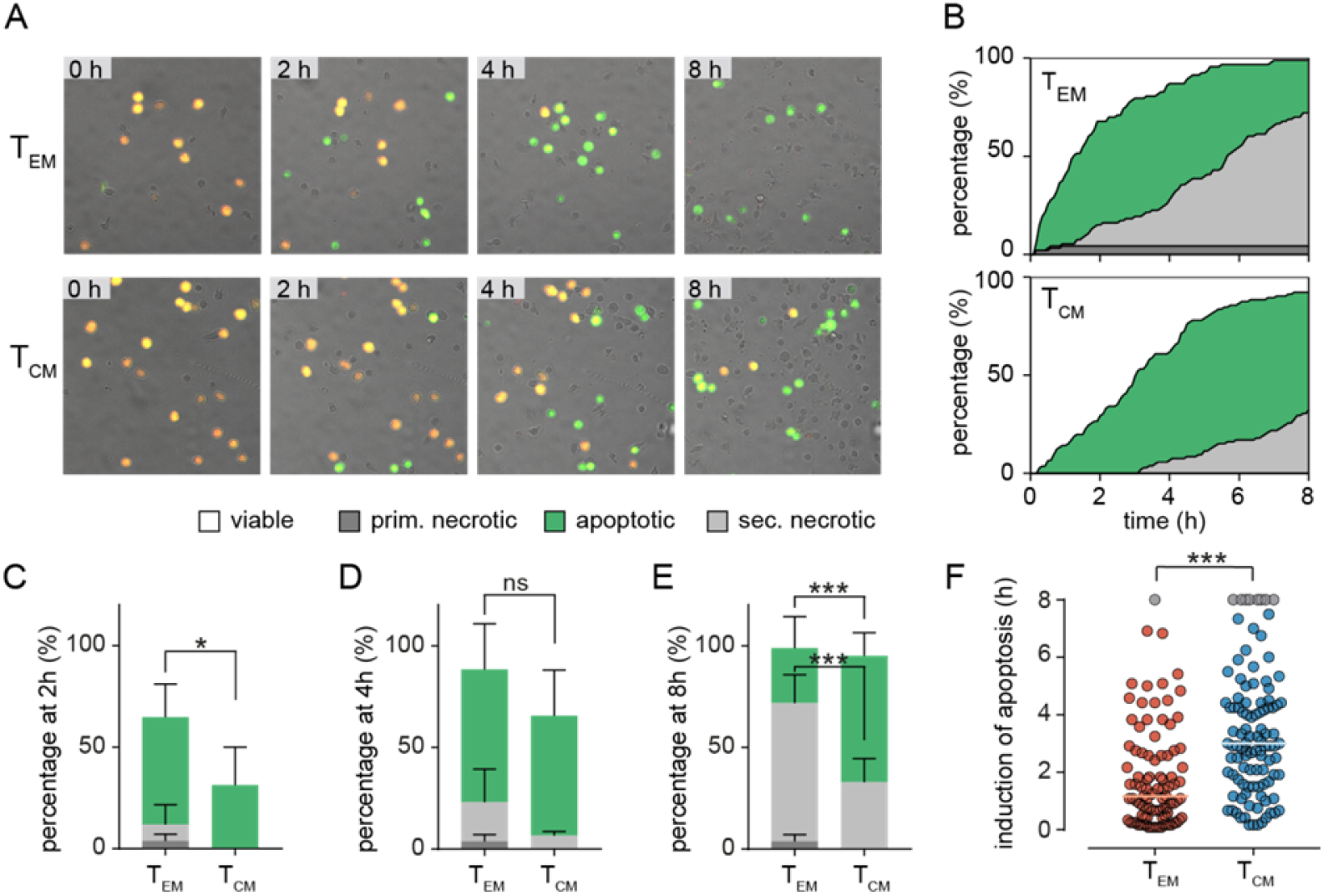
Single cell level T_EM_ and T_CM_ analyses reveal distinct cytotoxic efficiencies. A,B) Sorted TEM and TCM were co-incubated with SEA-pulsed NALM6 pCasper cells at an E:T ratio of 2:1 for 8h to analyze cytotoxicity on single cell level. A) Representative image section of T_EM_ or T_CM_ co-incubation. B) Quantification of target cell lysis events in a time resolved death plot showing the percentage of apoptotic target cells (green), primary necrotic target cell (dark grey), secondary necrotic target cell (light grey), necrotic/lysed (grey – for semi-automated analysis) or viable cells (white) by a manual. C)-E) Quantification of target cell lysis at 2 (C), 4 (D) or 8 (E) hours by manual analysis. F) Single cell values of duration/time of apoptosis for each analyzed target cell. Cells showing no signs of caspase activity at 8 hours are shown in grey. Each point reflects the Δt of apoptosis induction from a single target cell. Black bars show the median. N=4 donors, T_EM_ 94 cells, T_CM_ 105 cells. Statistical analysis was done using a two-way ANOVA. The orange and blue lines represent the median.

### Dissecting killing mechanisms of T_EM_ and T_CM_

Differences of killing efficiencies between T_EM_ and T_CM_ against target cells could be caused by different mechanisms including, among others, CTL migration speed or persistence, search strategies or different cytotoxic efficiencies during killer cell-target cell contacts. Considering that perforin, granzyme B and FasL (but not TRAIL) are expressed at a higher levels in T_EM_ than T_CM_, we first tested whether the different T_EM_ and T_CM_ killing efficiencies are caused by different efficiencies of the cytotoxic pathways.

Compared to control conditions (Fig. 6A, same plots as in 5B), anti-FasL blocking antibody had no effect on the induction of apoptosis and subsequent secondary necrosis by T_EM_ or T_CM_ mediated cell lysis (Fig. 6B) as also evident from the quantification of apoptosis and necrosis at 2, 4, and 8 hours for T_EM_ (Fig. 6E) and T_CM_ (Fig. 6F). One could argue that the antibody does not work, but we have shown previously that it works in NK cells and it also works in CTL if lytic granule exocytosis is blocked (see below).

**Figure 6:**
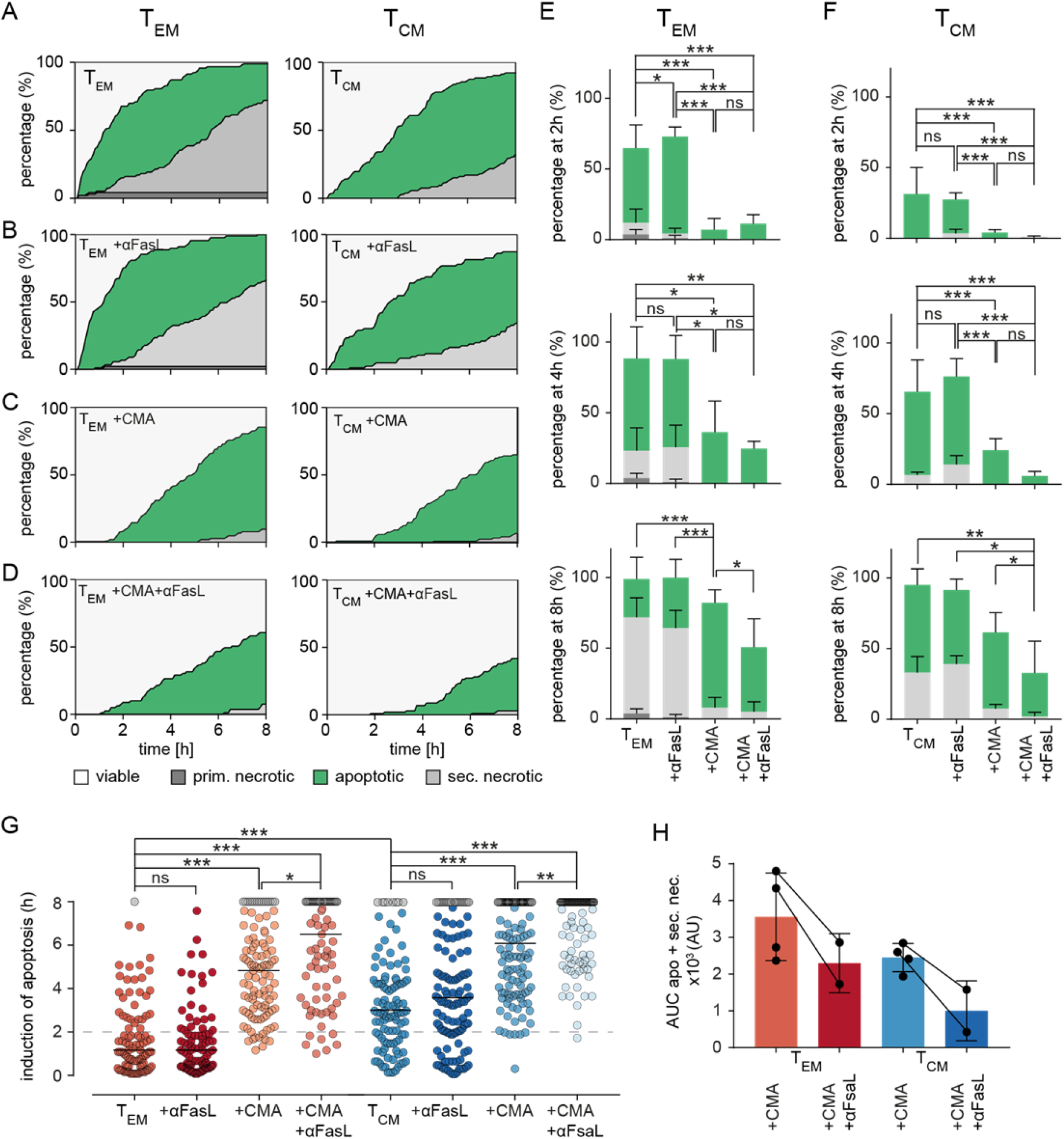
Dissecting killing mechanisms of T_EM_ and T_CM_. A) To analyze target cell lysis by perforin-mediated or FasR-mediated killing mechanisms, sorted T_EM_ or T_CM_ and target cell were either un-treated as control (A) or treated with inhibiting FasL antibodies NOK-1 and NOK-2 (10 μg/each) (B), with 50 nM CMA (C) and with inhibiting FasL antibodies and CMA in combination (D). Effects of treatments on T_EM_ and T_CM_ are shown as representative death plots, respectively (A-D). E,F) Quantification of target cell lysis at 2h, 4h and 8h for T_EM_ (E) or T_CM_ (F). G) Δt of apoptosis for each analyzed target cell lysed by treated T_EM_ or T_CM_ in comparison to untreated (T_EM_ or T_CM_) control cells. H) Impact of FasL blocking quantified by analysis of area under the curve (AUC). Each point reflects the Δt of apoptosis induction from a single target cell. n=4 donors, T_EM_: 94 cells, +αFasL: 89 cells, +CMA: 103, +CMA+αFasL (n=2 donors): 79, T_CM_: 105 cells, +αFasL: 122 cells, +CMA (n=2 donors): 122 cells, +CMA+αFasL: 102 cells. Statistical analysis was done using a two-way ANOVA.

The release of perforin from lytic granules can be blocked very well by low concentrations of concanamycin A (CMA) ((18)). CMA clearly reduced target cell apoptosis and it almost eliminated primary and secondary necrosis in both T_EM_ (Fig. 6C, quantification in Fig. 6E) and T_CM_ (Fig. 6C, quantification in Fig. 6F). We conclude that perforin is required for the fast apoptotic killing by T_EM_ and also for the slightly slower one of T_CM_ and its absence wipes out necrosis. Without perforin the big difference in cytotoxic efficiency between T_EM_ and T_CM_ is diminished. Thus, the higher perforin content of T_EM_ compared to T_CM_ is involved in their increased cytotoxic efficiency.

The inhibition of perforin release by CMA in combination with anti-FasL blocking antibody significantly reduced the percentage of apoptotic target cells in the late phase of the measurement for both, T_EM_ and T_CM_ (slightly stronger in T_CM_) (Fig. 6D-F).

To quantify the exact time points of apoptosis induction in the target cells, every individual cell was tracked over time. The exact time point of apoptosis could be quantified by loss of FRET-signals and an increase of GFP intensities (causing a switch from orange to green fluorescence). Quantification of 89 (T_EM_) or 122 (T_CM_) cells treated with CMA drastically prolonged the induction of apoptosis and as expected, even more dramatically in T_EM_ (by median increased by 220 min) compared to T_CM_ (median increased by 185 min) (Fig. 6G). Inhibition of the Fas/FasL pathway with anti-FasL blocking antibody again indicated no significant difference compared to control conditions in T_EM_ and T_CM_ subtypes. However, the combined treatment with CMA and anti-FasL blocking antibody delayed the induction of apoptosis further but to a very similar value for both subtypes, 100 min for T_EM_ and 115 min for T_CM_, respectively. Exploring the area under the curve (AUC) of the apoptotic and secondary necrotic events occurring over the whole experiment, allows the visualization and quantification of FasL-FasR interaction induced cytotoxicity (Fig. 6H). The additional inhibition of FasL, decreases the AUC of combined apoptotic and secondary necrotic cells for T_EM_ by 35.3% and for T_CM_ by 59.2% compared to CMA alone.

Taken together these data indicate a minor role of Fas-mediated target cell lysis for both subtypes, if perforin-mediated target lysis is functional. After blocking perforin function and subsequent induction of apoptosis by granzyme B, apoptotic target cell death induced through the FasL-FasR interaction is increasingly important for target cell lysis. Perforin-mediated target cell lysis appears to be prominent in both subsets, but the compensatory role of FasL-mediated target cell lysis appears to have a higher impact in T_CM_ compared to T_EM_.

Taken together this means that complete inhibition of perforin release by CMA and (at least) partial block of FasL-induced target cell death made both T_EM_ and T_CM_ less efficient serial killers. Nevertheless, T_EM_ were still slightly more efficient than T_CM_.

Thus, there may be additional factors guiding T_EM_ efficiency in comparison to T_CM_. To test this, we quantified T_EM_ and T_CM_ target cell killing in detail. First, we quantified how long T_EM_ or T_CM_ needed to find their first target cell (Fig. 7A). We did no distinguish whether this contact was successful or not, i.e. resulting in target cell destruction or not after T_EM_ or T_CM_ left the target. There was no difference between the time to first contact (Fig. 7A, median T_EM_ 42.5 min to T_CM_ 40 min; n=96, 106 cells, respectively), indicating that T_EM_ and T_CM_ migration patterns or search strategies do not influence the likelihood of finding a target cell.

**Figure 7:**
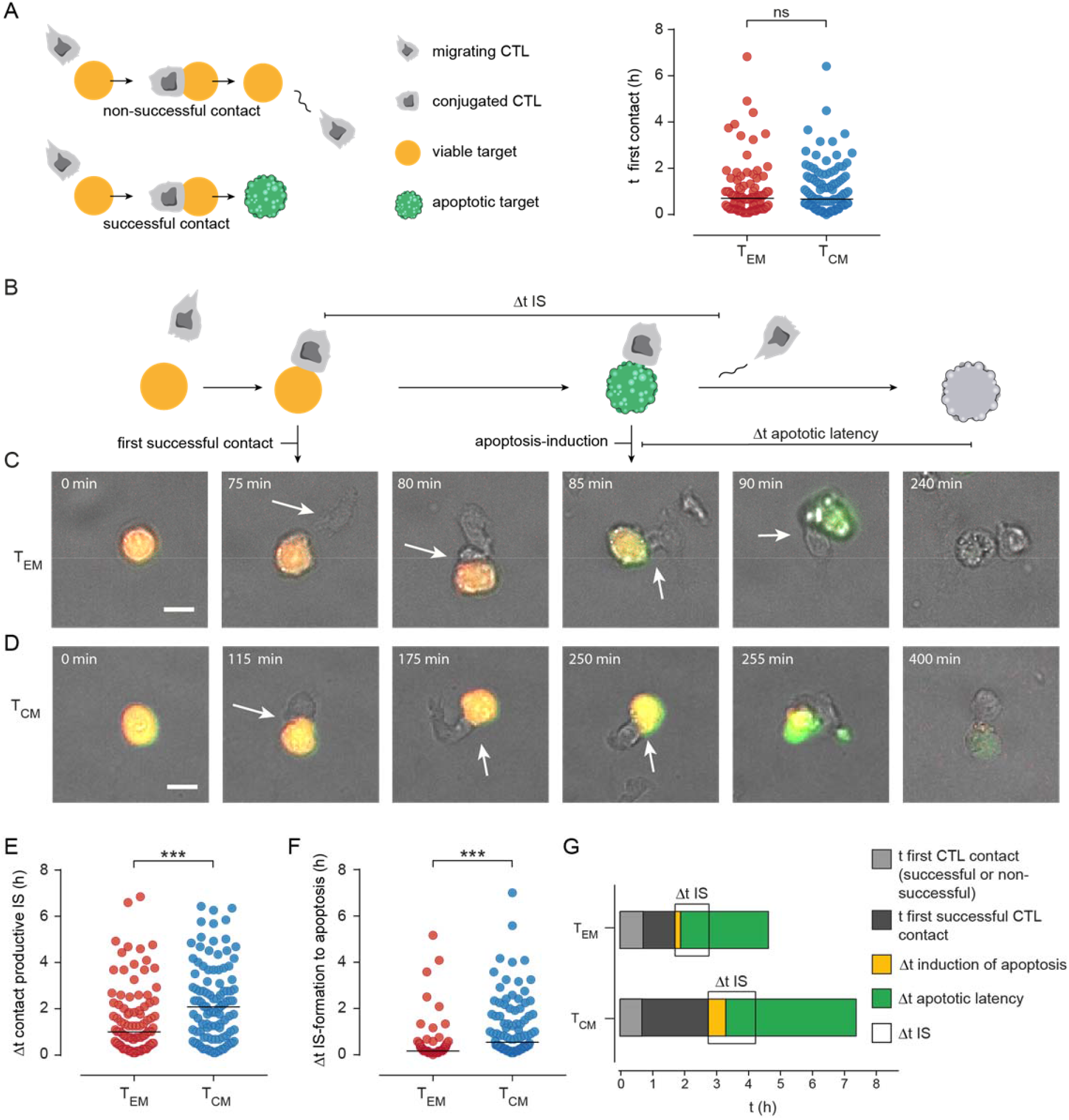
Manual tracking CTL-target cell contacs/interaction uncovers enhanced efficiency of immune synapse (IS) formation in T_EM_. A) Scheme of CTL-target cell interaction: When a CTL approaches a target cell this can result in a non-productive (no induction of apoptosis or necrosis, CTL leaves the viable target after minutes – hours) or in a productive contact that results in apoptosis or necrosis of the target cell. To check if differences in target cell lysis efficiency of T_EM_ and T_CM_ are dependent on migration the time point of the first contact (whether non-productive or productive) was compared. B) Scheme to illustrate the kinetics of productive CTL-target cell interaction: t of first productive contact (Δt of duration from start of the measurement until formation of a productive/lytic synapse. Time point of apoptosis induction (first signs of caspase activity detected by green fluorescence. Δt apoptotic latency: from induction of apoptosis until necrotic lysis of the cell (duration a cell was apoptotic) C and D) Representative images of T_EM_ (C) or T_CM_ (D) mediated target cell lysis. E) Single-cell data of time point of productive IS formation. F) Single-cell data of Δt of apoptosis induction (from IS-formation to first signs of caspase activity). G) Stacked bar graphs to summarize manually tracked single cell data from T_EM_ and T_CM_ mediated target cell lysis. t of first contact successful or non-successful (light grey). t of formation of the first productive contact (dark grey), Δt of apoptosis-induction (yellow), Δt of apoptotic latency (green). Δt IS: Duration of an IS between T_EM_ or T_CM_ and a target cell is shown in light grey. Durations of the phases are show as median. n=4 donors, T_EM_ 94 cells, T_CM_ 100 cells.

Next, we analyzed killing sequences in detail as displayed in Fig. 7B, examples for a T_EM_ and T_CM_ are shown in Fig. 7C, D. We quantified the time to the first successful contact, i.e. the time it took a T_EM_ or T_CM_ needed to contact a target which was killed by the CTL. The time to the first successful contact forming an IS was on average 1 hour for T_EM_ (Fig. 7C, statistics in 7E) and this was significantly shorter than the more than 2 hours it took T_CM_ (Fig. 7D, statistics in 7E).

In addition, during a successful response the time from IS formation to apoptosis induction was significantly shorter for T_EM_ than for T_CM_ (Fig. 7D, E, statistics in 7F, median 10 versus 55 min, p<0.0001) and IS duration was also shorter in case of T_EM_. The differences between T_EM_ and T_CM_ are summarized in Fig. 7G. Time to first successful contact (dark grey), duration of IS, time between IS formation and apoptosis induction in targets and also the time between apoptosis and secondary necrosis are all significantly shorter in T_EM_ than T_CM_.

In summary, whereas T_EM_ and T_CM_ need the same time to find a target cell T_CM_ need much longer than T_EM_ to establish the first contact with a subsequent successful killing. This means that T_CM_ have more failures, i.e. contacts with targets not resulting in killing. In addition, all measured time intervals (IS length, time to induce apoptosis, time to induce secondary necrosis) are shorter for T_EM_ compared to T_CM_.

## Discussion

Insights in single cell cytotoxic efficiencies of CTL subtypes are important to understand their function and to optimize adaptive T cell therapies. While the well-established and long-known ^51^Cr- release assay (24) is still considered the gold standard for detection of cytotoxicity in a CTL population, radioactivity and the lack of kinetic information are significant limitations. This led to the development of many alternative assays to quantify cytotoxicity (25–32). However, these are all population assays, they have no single cell resolution and do not discriminate between apoptosis and necrosis. Therefore, in this work a FRET-based caspase-dependent cytotoxicity assay was developed for CTL, which was previously shown to discriminate between apoptosis and necrosis at a single cell level (18, 33). The construct used here for quantification of caspase/protease is not the only one available, and others have been used to detect cell apoptosis by luciferase activity, quenching, or change in the fluorophore localization (20, 34, 35). However, none of these have been used for the detection of single CTL cytotoxicity. The FRET-based assay presented here works very well for CTL (this study) and NK cells (18).

This study combined population measurements and the single cell assay that allowed the precise quantification of T_EM_ and T_CM_ cytotoxic efficiencies. Both subtypes kill their targets mainly in a perforin-dependent manner but use also death receptors including FasR/FasL. The difference in perforin-dependent cytotoxicity, which is probably a result of the different perforin levels, is the major reason for the better cytotoxic efficiency of T_EM_ compared to T_CM_. In addition the lytic immunological synapse is more efficient for T_EM_-target cell contacts then for the ones with T_CM_. If this is also related to the perforin content is not clear at present. These data point however in a direction that CTL-target cell contact length is well controlled by a mechanism that measure the outcome of the lytic synapse.

The deep understanding of CD8^+^ T cell subtype development, composition and killing mechanisms are of great importance to optimize future adaptive cell therapies such as the use of CAR T cells. To date, molecular signatures, the signaling pathways of cytotoxic mechanisms and the cytotoxic receptor repertoire of death receptor or perforin-mediated target cell lysis of different CD8^+^ T cell subtypes have been comprehensively investigated, frequently flow cytometry or more recently mass cytometry based methods have been used (36–38). In contrast, the functional *in vitro* analysis of the cytotoxicity of human CTL subtypes is not on a comparable level, especially at the single cell level. In this manuscript we have elucidated the cytotoxic potential of human CD8^+^ T cell memory subtypes T_EM_ and T_CM_ and the relevance of their cytotoxic mechanisms. In our hands, the staphylococcal enterotoxin A (SEA)-stimulation of CD8^+^ T cells, which permits a subsequent analysis of the cytotoxicity against of SEA-loaded target cell (39–41), preferentially led to the differentiation into a TCM subtype. Nevertheless, the size of the T_EM_ and T_N_ populations was still large enough to have clearly defined subtypes available for further analysis. Only the terminally differentiated T_EMRA_ could not be included in further analysis as a limitation of SEA stimulation.

From a clinical perspective these data suggest a potential advantage of a selection of CTL subtypes for virus specific T cells, e.g. CMV-specific, or for the generation of CAR T cells. A limiting factor is certainly that patient-derived T cells are often poor in quantity and quality due to many previous therapies including purine analogues among others, so that a pre-selection of the initial T cell subset requires more efficient transduction and expansion protocols. However, superior results have been shown with CAR T cells in fixed CD4:CD8 ratios. A fixed composition of different T-cell subsets would be more feasible for off-the shelf products. On the other hand, different costimulating domains of CARs lead to different phenotypes of CAR T cells, i.e. CD19 scFv/41-BB-CD3ζ to T_CM_ phenotype and CD19 scFv/ CD28 CD3ζ to T_EM_ phenotype.

The data of the present manuscript suggest that CTL, and thus further developed CAR T cells have the ability to kill target cells harboring defective apoptosis mechanisms like del17p, by direct cytolysis, which is clinically very relevant. In the future, resistance mechanisms should be investigated, since CAR T cell failures have partially persistent CAR T cells and tumor cells partially continue to express CD19. The data of this work suggest that resistance mechanisms against perforin might be of particular interest in this context. To consider strengthening the effect of compensatory mechanisms such as the death domain ligands FasL and TRAIL therapeutically, for example by BCL2 inhibitors.

## Materials and methods

### Ethical approval

Research with human PBMC has been approved by the local ethic committee (84/15; Prof. Dr. Rettig-Stürmer). We got the leukocyte reduction system (LRS) chambers, a by-product of platelet collection from healthy blood donors, from the local blood bank (Institute of Clinical Hemostaseology and Transfusion Medicine, Saarland University Medical Center). All blood donors provided written consent to use their blood for research purposes.

### Reagents

Anti-human CD178 was obtained from BD Biosciences, anti-human CD95 Apo 1-1 was from Enzo Life Sciences. Flow Cytometry antibodies (for details see Flowcytometry) and apoptosis inducing TRAIL-Antibody were purchased from Biolegend. IL-7 was from Peprotech, IL-12 and IL-15 were from Miltenyi. Nucleofector Kit V was from Lonza. The original vector pCasper-GR (#FP971) was from Evrogen. BSA, fibronectin and staurosporine from Streptomyces sp. were from Sigma-Aldrich. The Dynabeads CD8 Positive Isolation Kit, fetal bovine serum (FBS) (EU-approved), IL-2, G418, PBS, RPMI-1640, AIM-V medium, penicillin/streptomycin (P/S) and were from ThermoFisher Scientific. Concanamycin A (CMA) was from Santa Cruz Biotechnology. All other chemicals and reagents not specifically mentioned were from VWR, ThermoFisher Scientific or Sigma-Aldrich.

### Cells lines and primary human CD8^+^ T cells

Raji cells (ATCC, #CCL-86) and NALM6 cells (DSMZ, #ACC128) were cultured in RPMI-1640 (ThermoFisher Scientific) supplemented with 10% FBS and 1% Penicillin/Streptomycin at 37°C and 5% CO_2_. Human peripheral blood mononuclear cells (PBMCs) were isolated from the leukoreduction system chamber (LRSC) as a by-product from healthy thrombocyte donation as described previously (42).

PBMCs were stimulated with staphylococcal enterotoxin A (SEA; 0.5 μg/ml) in a 24 well plate (density of 1–1.5 × 10^8^ cells/ml; 37°C for 1 h). PBMCs were resuspended at a density of 2–4×10^6^ cells/ml in AIM-V medium supplemented with 10% FBS and 100 U/ml recombinant human IL-2. 5 days after stimulation, SEA-specific CTL were positively isolated using a Dynabeads CD8 Positive Isolation Kit (ThermoFisher Scientific).

Purified CD8^+^ cells were cultured in AIMV medium supplemented with 10% FBS and 100 U/ml recombinant human IL-2 (ThermoFisher Scientific) and used for sorting of CD8^+^ T cell subsets and for further experiments three days after isolation.

### Generation of stable NALM6 pCasper cell line

The sequence encoding for Casper3-GR was amplified from pCasper3-GR (Evrogen # FP971) with the following primers introducing an XhoI recognition site at both ends of the amplicon. Forward primer: 5’-CTCGAGGCCACCATGGTGAGCGAG-3’, reverse primer: rev 5’-GACGAGCTGTACCGCTGACTCG AG-3’. The amplicon was subcloned into the XhoI site of a previously modified pGK-Puro-MO70 vector backbone were CITE-EGFP sequence was deleted (43). The generated plasmid encoded puromycin resistance gene and was used for selecting transfected NALM6 cells. 1 × 10^6^ NALM6 target cells were transfected with 1 μg plasmid using the Nucleofector 4D (SF cell line Kit, program CV-104, Lonza) following the manufacturer’s instruction. After 48 hours, selection started with 0.2 μl/ml puromycin (1 mg/ml stock solution). To enrich the pCasper GFP^+^/RFR^+^ population for clonal selection, GFP- and RFP-positive cells were sorted on a FACSARIAIII sorter and plated as 1 cell/per well in 96 well plates. GFP- and RFP-positive clones were selected by the ImageXpress Micro XLS screening image analysis system (Molecular Devices), expanded and frozen in 90%FCS/10%DMSO for further use.

### Flow Cytometry

The frequency of CD8^+^ T cell subpopulations was examined by intra- and extracellular stainings. 5×10^5^ cells were washed twice in PBS/0.5% BSA, stained with PerCP-conjugated anti-human CD3 (SK7, Biolegend), FITC-conjugated anti-human CD8 (SK1, Biolegend), PE/Cy7-conjugated anti-human CD45RA (Hl100), Biolegend) or CD45RO (UCHL-1, Biolegend), PerCp/Cy5.5-conjugated anti-human CD62L (DREG-56, Biolegend), Alexa Fluor-647 conjugated anti-human CD197 (150503, BD Biosciences) antibodies in PBS/0.5% BSA, washed twice and recorded on a BD FACSVerse flow cytometer (BD Biosciences). For intracellular stainings, cells were fixated in 4% PFA in PBS for 20 min, washed in PBS/0.5% BSA, permeabilized for 10 min in PBS/0.1% saponin and stained with FITC-conjugated anti-human Perforin (γG9, Biolegend) and PE-conjugated anti-human Perforin antibody (B-D 48, Biolegend) in 0.1% PBS/0.1% saponin. After staining cells were washed and acquired on a FACSVerse flow cytometer.

For analysis of degranulation and staining of the death receptors FasL and TRAIL, 3×10^5^ SEA-pulsed target cells or target cells without SEA as control were seeded in wells of a 96-well plate. 6×10^5^ SEA-CTL were added to each well and cells were co-incubated in the presence of Brilliant Violet 421-conjugated anti-human CD107a antibody. After 4 h cells were collected on ice, washed (4°C) and a surface staining of CD45RA, CCR7, PE-conjugated anti-human CD178 (NOK-1, Biolegend) or PE-conjugated anti-human CD253 (RIK-2, Biolegend) was performed, cells were washed before analysis on a FACSVerse flow cytometer (BD Biosciences).

T_EM_ or T_CM_ were generated from human PBMC (peripheral blood mononuclear cell) using a widely accepted protocol for polyclonal activation of T cells by staphylococcal enterotoxin A (SEA) (40, 44) that importantly allows the subsequent analysis of SEA-dependent cytotoxicity (30, 39). For sorting 4×10^7^ SEA-CTL were stained with anti-CD62L and anti-CD45RO in PBS + 0.5% BSA, washed, strained (30 μm) and sorted on a BD FACSAria III (70 μm nozzle). Cells were collected in PBS/0.5% BSA/25% FBS, washed and cultured in AIM-V + FBS + 100 ng/ml IL-2. T_EM_ received 25 ng/ml IL-2, 5 ng/ml IL-12, 10 ng/ml IL-15, T_CM_ received 25 ng/ml IL-2, 5 ng/ml IL-7 and 10 ng/ml IL-15. Cells were measured on a BD FACSVerse flow cytometer and data analysis was performed using FlowJo.

### Real-time killing assay

To quantify the cytotoxicity of SEA-stimulated CD8^+^ T cells against SEA-pulsed Raji or NALM6 target cells we used a time resolved, real-time killing assay that was carried out essentially as previously described (30). Briefly, target cells (Raji or NALM6) were pulsed with 1 μg/ml SEA in AIM-V medium at 37°C and 5% CO_2_ for 30 min. Next, cells were loaded with 500 nM Calcein-AM in AIM-V medium supplemented with 10 mM HEPES for 15 min at room temperature. After one washing step, 2.5 × 10^4^ target cells per well were pipetted into a 96-well black plate with clear-bottom (#7342480, VWR). After a short rest (15 min) to let target cells settle down, effector cells were added at the indicated effector to target ratio (E:T) and target cell lysis was measured in a Genios Pro (Tecan) reader using bottom reading function over 4 h at 37°C every 10 minutes. Maximal killing rates were calculated as the maximum increase between two subsequent measured points (10 minutes).

### 2D single cell imaging and target cell death analysis

Single cell killing assay measurements were run on an ImageXpress Micro XLS screening image analysis system (Molecular Devices). The detection of apoptotic and necrotic target cells with the apoptosis sensor Casper-GR construct was carried out as described before ((18)). Briefly, NALM6 pCasper target cells were pulsed with 1 μg/ml SEA in AIM-V medium at 37°C and 5% CO_2_ for 30 min, washed taken up in 100 μl of phenol-free RPMI-1640/10% FBS and seeded into fibronectin coated wells (coating 30 min, 50 μl of 0.1 mg/ml fibronectin per well) of a 96-well plate (1×10^4^ targets per well). After a short rest (30 min) 2×10^4^ effector cells in 100 μl of phenol-free RPMI-1640/10% FBS were added per well and the measurement was started. Over 8h bright field, GFP- and FRET-signals were acquired witch a 20× objective from 1-4 positions per well at a 5 min interval. Objects were excited via Spectra X LED illumination (Lumencor) using LED 470/24 for GFP. The filter sets for FRET were 472/30 nm for excitation and 520/35 nm for emission to measure GFP and 641/75 nm to measure RFP. A Nikon Super Fluor objective (20×/NA 0.75) was used.

### Treatment with inhibitory substances

To inhibit perforin-mediated target cell lysis effector cells were preincubated for 2 h in phenol-free RPMI-1640/10% FBS supplemented with 50 nM concanamycin A (CMA) and 50 nM CMA was also present during the measurements. 1×10^4^ targets per well were re-suspended in 100 μl phenol-free RPMI-1640/10% FBS/50 nM CMA. Effector cells were added in 100 μl of phenol-free RPMI-1640/10% FBS/50 nM CMA after preincubation.

To inhibit granzyme B-mediated target cell lysis, effector cells were preincubated for 2 h in phenol-free RPMI-1640/10% FBS supplemented with 10 μM z-AAD-CMK and 10 μM z-AAD-CMK was also present during the measurement. To this end 1×10^4^ targets per well were submitted in 100 μl of phenol-free RPMI-1640/10% FBS/10 μM z-AAD-CMK. Effector cells were added in 100 μl of phenol-free RPMI-1640/10% FBS/10 μM z-AAD-CMK after preincubation.

To block FasL- and TRAIL-mediated target cell lysis inhibitory anti-human antibodies were used to block the interaction of FasL or TRAIL with their respective apoptosis inducing receptors. To this end 1×10^4^ targets per well and 2×10^4^ effector cells per well were each preincubated for 2 h in 10 μg/ml anti-human CD178 antibodies (NOK-1 and NOK-2, BD Biosciences) or 5 μg/ml anti-human TRAIL antibody (RIK-2, Biolegend). After 2 h of preincubation effector cells were added to target cells.

### Image analysis

Images were analyzed as described in (18). Briefly, background correction via rolling ball, F/F0 normalization and donor ratio calculation were performed in ImageJ (NIH, version1.51d) and Microsoft Excel (Microsoft Office 2016). Following rolling ball background subtraction in ImageJ measurements were tracked manually with Speckle TrackerJ plugin for ImageJ.

### Manual target cell tracking and cell death analysis

In principle the manual analysis was carried out as described before (18). Briefly: Roughly 25 target cells were chosen randomly for each recorded position and tracked with Speckle Tracker J. The center of each target cell was marked at each time point manually. X,y positions and GFP- and FRET fluorescence signals were exported for every tracked cell with the speckle tracker intensity trajectories plugin. Data were imported into Excel to perform normalization to the starting value of a target cell and calculate the donor ratio for each cell. The GFP- and FRET-signal as well as the donor ratio were analyzed to determine the fate of a cell, viable (donor ratio > 0.75), apoptotic (donor ratio < 0.75) or necrotic (loss of both fluorescence signals, threshold FRET-value 10% above background signal; FRET< 150 units). This was used to identify the status of each tracked target cell at every time point. A cell being apoptotic prior to necrosis was defined as secondary necrosis. A necrotic event without a rise of the GFP-signal and an accompanying drop of the donor ratio was defined as primary necrosis. The sum of all detected cells per condition was set to 100% and the proportion of apoptotic, primary necrotic or secondary necrotic cells at any given time point was calculated and displayed in a color-coded death plot over time.

### RNA isolation and quantitative RT-PCR

Molecular biology was done as described before (45). Total RNA was isolated from 1.0 to 1.5 × 10^6^ CD8^+^ T-cells using TRIzol reagent (ThermoFisher Scientific) including 1 μl Glycogen (5 μg/μl, ThermoFisher Scientific). 0.8 μg total RNA was reverse transcribed to cDNA and 0.5 μl of cDNA was used for quantitative real time polymerase chain reaction (qRT-PCR). QRT-PCR was carried out in a CFX96™ Real-Time SystemC1000™Thermal Cycler (Biorad CFXManager). Primers used in this study: RNAPol and TBP as reference genes (45), FASL: forw 5′ GCACACAGC ATCATCTTT 3′, rev 5′ CAAGATTGACCCCGGAAGTA 3′; perforin (NM_005041 and NM_001083116) forw 5′ ACTCACAGGCAGCCAACTTT 3′, rev 5′ CTCTTGAAGTCAGGGTGCAG 3′; granzyme B, forw 5′ GAGACGACTTCGTGCTGACA 3′ and rev 5′ TCTGGGCCTTGTTGCTAGGTA 3′. QuantiTect primers for IFNγ, TNFα and TRAIL were purchased from Qiagen (QT00000525, QT00029162, QT00079212).

### Data and statistical analysis

Data were tested for significance using one-way Anova, Mann-Whitney-Test, Kruskall-Wallis-Test with a Dunn’s multiple comparisons post-test: * p<0.05; ** p<0.01; *** p<0.001; ns, no significant difference as stated in the figure legends. Data analyses were performed using Microsoft Excel 2016, FlowJo, Imaris (Bitplane, V8.1.2), ImageJ, and GraphPad Prism 7 software.

## Abbreviations

CD8^+^: effector memory (T_EM_)
CD8^+^: central memory (T_CM_)
CD8^+^: T cells (CTL)
LRS: (leukocyte reduction system) chambers

## Acknowledgements

We are very grateful to Prof. Hermann Eichler and the Institute of Clinical Hemostaseology and Transfusion Medicine at Saarland University Medical Center for obtaining human blood cells. We also are very thankful all human blood donors. We are very grateful to Carsten Kummerow and Christian Backes for introducing the Casper-GR system for single cell analysis, support with the data analysis and detailed discussions. We thank Susanne Renno for help with generation of stable cells lines and Sandra Janku for language proof reading. We thank Cora Hoxha and Carmen Hässig for cell preparation and excellent technical support. We are grateful to Elmar Krause and Jens Rettig for allowing us to use the flow cytometric sorting facility. This work was supported by the Deutsche Forschungsgemeinschaft (DFG, the collaborative research centers SFB 1027 (project A2 to MH), SFB 894 (project A1 to MH)) and by the Bundesministerium für Bildung und Forschung (BMBF, grant 031LO133).

## Author contributions

AK carried out the experiments, performed the analysis and was involved in writing the manuscript. GS and DA performed experiments. LT helped with writing the manuscript. MH contributed in planning the experiments and in writing the manuscript. ECS supervised the project and wrote the manuscript.

