## Supplementary Figure 1 for "Cytotoxic efficiency of human CD8^+^ T cell memory subtypes"

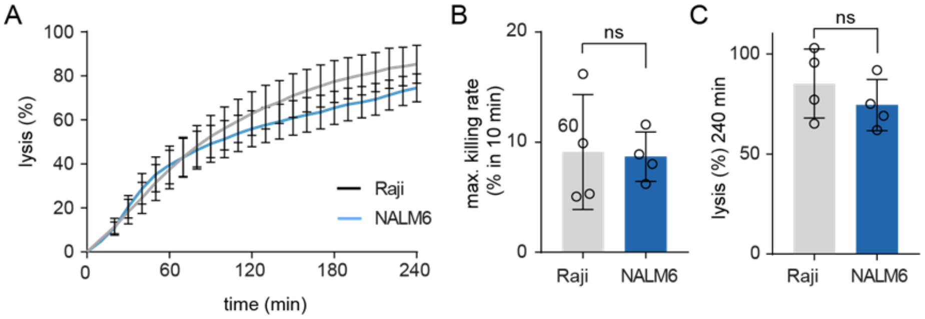


**Supplementary Figure 1: Comparison of Raji- and NALM6 target cell systems for SEA-mediated cytotoxicity.** A) Cytotoxicity of SEA-CTL against target cells Raji (grey) or NALM6 (blue) was analyzed by calcein-based real-time killing assay over 240 min. Effector to target ratio (E:T) was, n=4 donors, data are shown as mean +SD. Analysis of the maximal killing rate (B) and end point target lysis after 240 min (C).
